# Dispersal limitations of early life stages and sibling aggregations in a broadcasting *Millepora* hydrocoral, as revealed by parentage analysis

**DOI:** 10.1101/413963

**Authors:** Caroline E. Dubé, Emilie Boissin, Alexandre Mercière, Serge Planes

## Abstract

Dispersal is a critical process for the persistence and productivity of marine populations. For many reef species, there is increasing evidence that local demography and self-recruitment have major consequences on their genetic diversity and adaptation to environmental change. Yet empirical data of dispersal patterns in reef-building species remain scarce. Here, we document the first genetic estimates of local dispersal and self-recruitment in a broadcasting reef-builder, the hydrocoral *Millepora platyphylla*. Using microsatellite markers, we gathered genotypic information from 3160 georeferenced colonies collected over 9000 m^2^ of reef in three adjacent habitats in Moorea, French Polynesia; the mid slope, upper slope and back reef. Our parentage analysis revealed a predominance of self-recruitment with 58% of sexual propagules produced locally. Sexual propagules often settled at less than 10 meters from their parents and dispersal events decrease with increasing geographic distance. Limited dispersal among adjacent habitats via cross-reef transport was also detected. Sibship analysis showed that both full and half siblings recruit together on the reef, resulting in sibling aggregations. The identification of local families revealed discrepancies between dispersal patterns of sexual and asexual propagules. Self-recruits are dispersed with along-reef currents and settled in alignment with the location of their parents, while the dispersal of asexual fragments is heavily influenced by wave-driven cross-reef currents. Our findings highlight the importance of self-recruitment together with clonality in stabilising population dynamics, as it can enhance local sustainability and resilience to disturbance, but also raise uncertainties on the widely accepted high dispersal ability of broadcasting reef species.

## Introduction

Understanding patterns of dispersal is a major goal in ecology and conservation biology (Botsford et al., 2009; Cowen et al., 2007; Warner & Cowen, 2002). These patterns shape species’ distribution and abundance (Strathmann et al., 2002) and have major consequences for the persistence and adaptation of their populations (Garant et al., 2007; Gilmour et al., 2013; Underwood et al., 2009). For most marine species whose adults are sessile or relatively sedentary, the early life history includes a propagule stage that represents the first step for successful recruitment. Propagule dispersal depends on many biological and physical processes, including the survival and development rates of propagules (Doropoulos et al., 2016; Figueiredo et al., 2013; Johnson et al., 2014), larval behaviour (Gerlach et al., 2009; Paris et al., 2007), and hydrodynamic regimes and seascapes (Cowen et al., 2006; Cowen & Sponaugle, 2009; White et al., 2010).

In coral reefs, many organisms, such as fishes and scleractinian corals, rely on the dispersal of a larval stage for population replenishment and colonisation of fragmented habitats. Determining whether the maintenance of reef populations is more heavily influenced by self-recruitment (recruitment into a population from itself) or gene flow (longer-distance dispersal and connectivity) is an issue of much debate (Cowen et al., 2000; Strathmann et al., 2002). Genetic parentage data are an increasingly common source of empirical dispersal information, especially for reef fishes (Abesamis et al., 2017; Almany et al., 2017; Planes et al., 2009; Salle et al., 2016). For instance, parentage analysis has uncovered high levels of recruitment back to the parental source for some reef fish species (e.g., 30–60%, see Almany et al., 2007; Jones et al., 2005; Salles et al., 2016). This research reinforces the idea that reef populations are less open than previously thought.

In scleractinian corals, the extent of dispersal is largely governed by their reproductive biology and early life history ecology. Our ability to make inferences on what specific biological processes are driving dispersal and subsequent recruitment patterns in reef corals is limited due to their diverse dispersal strategies; gamete broadcasting, larval brooding and asexual reproduction pathways, which includes fragmentation, budding, polyp bail-out, asexually produced planula and embryo breakage (Harrison, 2011; Heyward & Negri, 2012). Nevertheless, field surveys and population genetic studies have suggested high levels of self-recruitment and limited dispersal in corals at some reefs (Baums et al., 2005; Concepcion et al., 2014; Gilmour et al., 2009; Shinzato et al., 2015; Torda et al., 2013). Recently, the application of genetic parentage analysis in reef corals has provided the first estimates of restricted larval dispersal in brooding species (Warner et al., 2016). Although broadcast spawning of gametes with planktonic development of larvae is the most common reproductive strategy in reef corals (Baird et al., 2009), dispersal patterns of their sexual propagules using parentage analysis have yet to be determined. Such information is essential to disentangle whether or not this reproductive strategy optimises larval dispersal as commonly thought.

*Millepora* hydrocorals, also called fire corals, are an important component of reef communities where they, similar to scleractinian corals, contribute to the accretion of reefs (Lewis, 2006; Nagelkerken & Nagelkerken, 2004). *Millepora* species inhabit a wide range of habitats (Dubé et al., 2017a; Lewis, 2006) and often grow into large colonies that pre-empt space and compete with scleractinian corals (Andréfouët et al., 2014; Dubé et al., 2016). Despite their importance in reef community dynamics, fire corals have been relatively understudied and not much is known with respect to their reproduction and dispersal patterns. *Millepora* species are broadcast spawners that alternatively reproduce by shifting from sexual reproduction to asexual pathway of fragmentation (Bourmaud et al., 2013; Lewis, 2006). The use of sexual/asexual modes of reproduction has been recorded in *Millepora platyphylla* at Moorea (Society Archipelago, French Polynesia), where habitat specific environmental conditions are thought to determine the levels of clonality (Dubé et al., 2017b). The spatial distribution of clone mates on a barrier reef (as in Moorea) has demonstrated that the dispersal of asexual fragments in fire corals was heavily influenced by an across-reef gradient of wave energy from offshore reefs towards lagoonal habitats (Dubé et al., 2017b). However, it remains unknown whether the dispersal of their sexually produced propagules is driven by cross-reef transport and dispersal among adjacent reef habitats. Although such information is crucial for understanding the replenishment and recovery of marine populations, there is no genetic study that has identified patterns of sexual dispersal and recruitment in these important reef-building species.

Here, we conducted an extensive field survey of 3160 georeferenced colonies of *M. platyphylla* over 9000 m^2^ of reef, based on replicated transects across three adjacent habitats (mid slope, upper slope and back reef) in Moorea. Using parentage analysis, i) we established the relative contribution of self-recruitment *versus* gene flow in the local replenishment of *M. platyphylla* population, ii) investigated dispersal patterns of sexual propagules within and across reef habitats, iii) determined how far siblings settle from each other and from their parents, and iv) assessed how fragmentation influences local recruitment in a partially clonal reef-building coral.

## Materials and Methods

### Model species

*Millepora platyphylla* is a gonochoric broadcast spawner that reproduces sexually by producing medusoids (modified medusa) and planula larvae (Bourmaud et al., 2013; Lewis, 2006). Medusoids are developed at the surface of the polyp coenosteum in cavities called ampullae and undergo sexual reproduction (Lewis, 1991). The medusoids are shed freely in the water and release their gametes at the surface in one hour post-spawning (short-lived). After fertilisation, the zooxanthellate planula larvae sink and move epibenthically (crawling not swimming) on the reef substratum before metamorphosing into a new calcifying polyp in one day to several weeks after spawning (Bourmaud et al., 2013). Fire corals also rely heavily on asexual reproduction through fragmentation for local replenishment (Dubé et al., 2017b), while the production of asexual larvae has never been documented so far within the *Millepora* genus.

### Population sampling and microsatellite genotyping

Between May and September 2013, a series of surveys were conducted on the north shore of Moorea, French Polynesia, across three adjacent reef habitats at Papetoai: two on the fore reef, the mid slope (13 m depth) and upper slope (6 m depth), and the back reef (< 1 m depth) (Fig. 1). Within each habitat, three 300 m long by 10 m wide belt transects were laid over the reef parallel to shore for a total of 9000 m^2^ of reef area surveyed. All colonies of *M. platyphylla* that were at least 50% within the transect borders were georeferenced by determining their position along the transect-line (0 to 300 m) and straight-line distance from both sides of the transect (0 to 10 m). From these measures, each colony was mapped with x and y coordinates. A map of the locations of each colony was produced using R (R Development Core Team, 2016). The colony size (projected surface) of each colony was estimated (in cm^2^) from 2D photographs using ImageJ 1.4f (Abràmoff et al., 2004). To determine the gender in *Millepora* it requires the presence of medusoids at the surface of the colony as well as their release in the water column to observe the reproductive gametes (oocytes and sperm sacs). Such observations are difficult in the field and have only been made from specimens in aquariums so far (Bourmaud et al., 2013; Lewis, 1991; Soong & Cho, 1998; Weerdt, 1984). Consequently, the sex of the colonies was not determined for this study. Small fragments of tissue-covered skeleton (< 2 cm^3^) were sampled during field surveys and preserved in 80% ethanol and stored at –20°C until DNA extraction.

**Figure 1.**
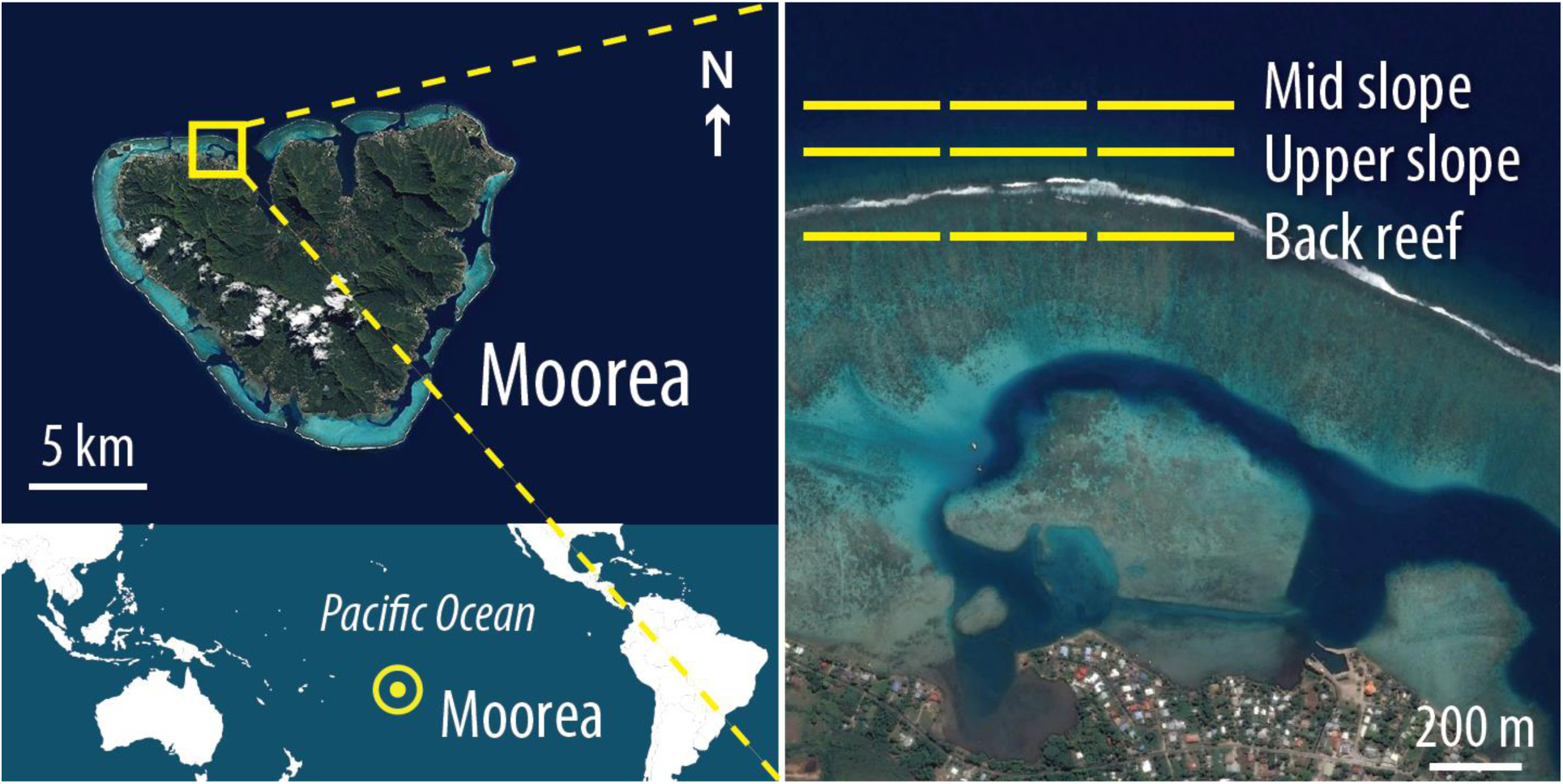
Aerial views showing the study area in Moorea, French Polynesia, and the locations of the three belt transects (300 x 10 m) within the three surveyed habitats. The world map was obtained from Aix-Marseille University (http://www.d-maps.com) and images from Google Earth (Map data © 2015 Google, DigitalGlobe). The figure was created using Adobe Photoshop CS6 software.

All samples were incubated at 55 °C for 1 hour in 450 µL of digest buffer with proteinase K (QIAGEN, Hilden, Germany) and DNA was extracted using a QIAxtractor automated genomic DNA extraction instrument, according to manufacturer’s instructions. Each colony was amplified at twelve polymorphic microsatellite loci in four multiplex polymerase chain reactions (PCRs) using the Qiagen Multiplex PCR Kit (Table S1). Further details on these loci and the genotyping procedure are described in Dubé et al. (2017c). Samples were sent to GenoScreen platform (Lille, France) for fragment analysis on an Applied Biosystems 3730 Sequencer with the GeneScan 500 LIZ size standard. All alleles were scored and checked manually using GENEMAPPER version 4.0 (Applied Biosystems, Foster City, CA).

### Multilocus genotypes and summary statistics

Multilocus genotypes (MLGs) were identified in GENCLONE version 2.0 (Arnaud-Haond & Belkir, 2007). To assess the discriminative power of the microsatellite markers, we estimated the genotype probability (GP) for each locus and a combination of all loci in GENALEX version 6.5 (Peakall & Smouse, 2006). Colonies (ramets) with the same alleles at all loci were assigned to the same MLG (genet) and considered to be a product of asexual reproduction when GP < 0.001. To evaluate the probability that two identical MLGs arise from distinct random reproductive events, the probability of identity, *P*_(ID)_, was estimated (Waits et al., 2001). The probability of exclusion (*P*_EX_) was also computed in GENALEX and indicates the efficiency of a panel of microsatellite markers to exclude unrelated individuals when both parents are unknown (Jamieson & Taylor, 1997). Population genetic analyses were performed after the removal of all clonal replicates (genet level). Indices of genetic diversity, including the total number of alleles per locus (N_A_), observed (*H*_O_) and expected (*H*_E_) heterozygosity (Weir & Cockerham, 1984) were estimated in GENALEX. Deviations from HWE (*F*_IS_) were assessed with a permutation procedure (N = 1000) implemented in GENETIX version 4.02 (Belkhir et al., 1996). Pairwise relatedness coefficient (Lynch & Ritland, 1999) and pairwise geographic distances between colonies were computed in GENALEX.

### Parentage analyses

Kinship analyses were performed in COLONY version 2.0 (Jones & Wang, 2010) to infer sibship and parentage relationships using individual MGLs. For the following analyses, only mature colonies (> 20 cm^2^) were considered as potential parents. Colonies smaller than 20 cm^2^ were considered as juveniles (i.e. non reproductive colonies, a threshold currently used for scleractinian species, see Penin et al., 2010; Sandin et al., 2008) and assumed to be the pool of potential offspring. Parent-offspring relationships were assessed with all parental genotypes entered into the analysis as candidate fathers because the sex of colonies could not be determined. COLONY was also used to identify siblings (full- and half-sibs) in the juvenile samples only. COLONY was launched with the following parameters for each of the three medium length runs: both sexes are polygamous and the organism is dioecious, with inbreeding, full-likelihood method and a medium likelihood precision. Each run included explicit marker error rates, allele frequencies for the studied population computed with GENALEX and 80% sampled candidate fathers and unknown maternal sibs. Only inferred assignments with a probability of 0.95 or greater were considered for the results. Pairwise geographic distances for all parentage relationships (parent-offspring, parent-parent, full and half siblings) were estimated in GENALEX.

### Statistical analyses

Two-tailed binomial tests (proportion test) were used to assess differences in proportion of juveniles, as well as variations in the contribution from self-recruitment, among the three surveyed habitats. Pearson’s correlation coefficient was used to determine whether the proportion of dispersal events (parent-offspring, full- and half-sib pairs) decreases with increasing geographic distance, while differences for proportions among geographic distances were tested using two-tailed proportion tests. Chi square tests with Monte Carlo simulation (1000 replicates) and Bonferroni corrections (to compensate for multiple comparisons) were used to assess for differences in dispersal patterns among full and half siblings. Pearson’s correlation coefficient was also used to determine whether the number of offspring produced by each of the identified parents increases with their number of ramets (parents with or without clones of the same genotype) and colony size. Furthermore, differences in dispersal distances between offspring assigned to parents belonging to a clonal genotype (parents with clones) *versus* non-clonal genotype (parents without clone) were assessed using a two-tailed binomial test (Student’s t-test). All statistical analyses were performed in the R programming environment version 3.1.2 (R Development Core Team, 2016).

## Results

### Population sampling, clonality and genetic diversity

The population density was approximately one colony per 10 m^2^ and the mean pairwise geographic distance between all colonies within the entire study area was 347 m (± 228 SE; range: 0.01–979 m; median = 310 m). The size-frequency distribution of the population was skewed towards small colonies (g_1_ = 0.395; Pnorm < 0.001) with 64% of all colonies below 125 cm^2^ (Fig. S1). Of all surveyed colonies, 1059 were considered as juveniles (< 20 cm^2^) and 2101 as adults (Fig. 2 and Table 1).

**Table 1.**
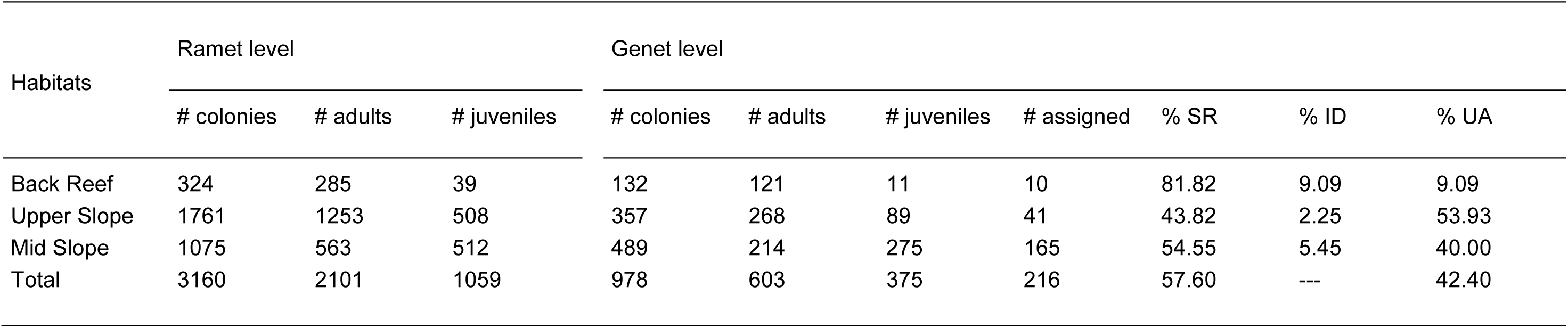
Details of *Millepora platyphylla* population sampling based on colony size (ramet level) and parentage assignments across the three surveyed habitats at Moorea. Percentages give the proportions of juveniles within that habitat that were i) offspring of parents sampled at the same habitat (SR = self-recruitment), ii) offspring of parents sampled at other habitats (ID = inter-habitat dispersal), or iii) were not assigned to parents sampled within the entire area surveyed (UA = unassigned).

| Habitats | Ramet level |  |  | Genet level |  |  |  |  |  |  |
| --- | --- | --- | --- | --- | --- | --- | --- | --- | --- | --- |
|  | # colonies | # adults | # juveniles | # colonies | # adults | # juveniles | # assigned | % SR | % ID | % UA |
| Back Reef | 324 | 285 | 39 | 132 | 121 | 11 | 10 | 81.82 | 9.09 | 9.09 |
| Upper Slope | 1761 | 1253 | 508 | 357 | 268 | 89 | 41 | 43.82 | 2.25 | 53.93 |
| Mid Slope | 1075 | 563 | 512 | 489 | 214 | 275 | 165 | 54.55 | 5.45 | 40.00 |
| Total | 3160 | 2101 | 1059 | 978 | 603 | 375 | 216 | 57.60 | --- | 42.40 |

**Figure 2.**
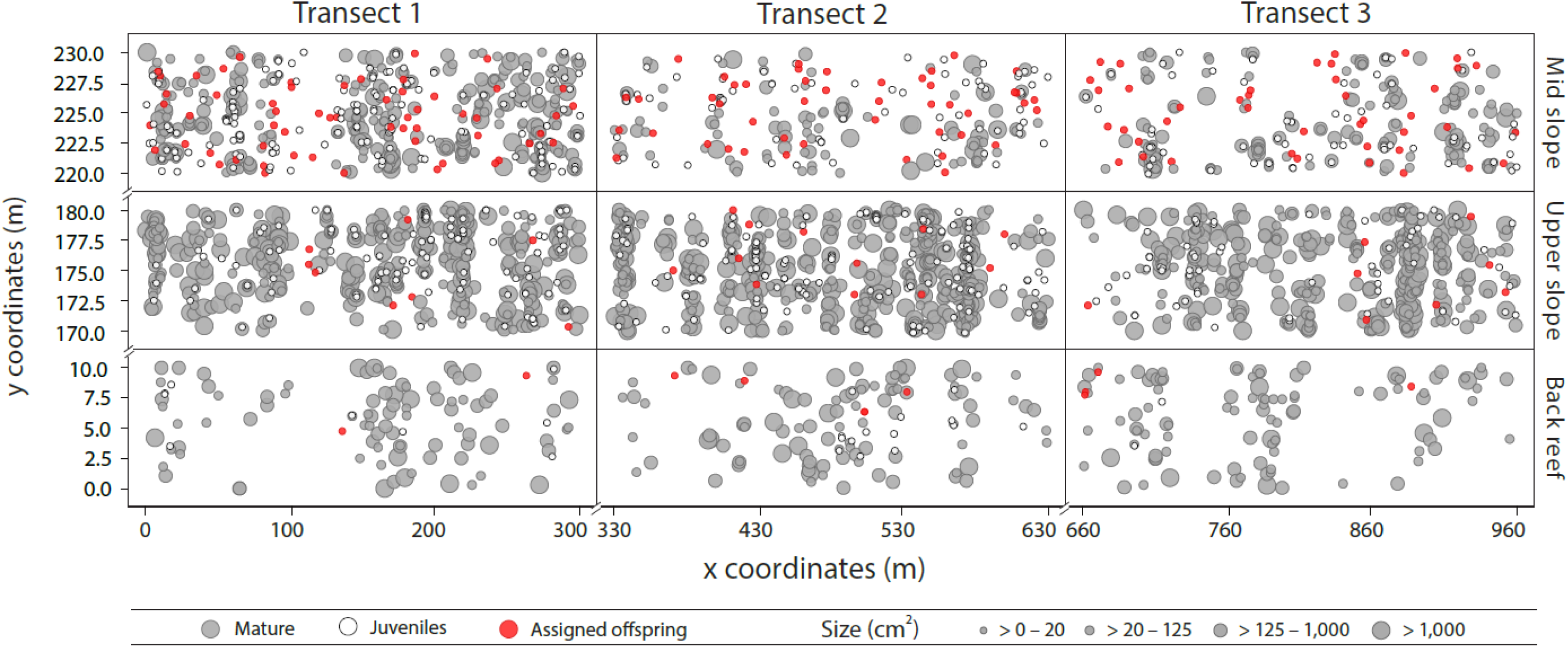
Spatial distribution and size of the 3160 colonies sampled across the three surveyed habitats within three belt transects (300 x 10 m). Adult colonies (potential parents, N = 2101) are shown in grey and juvenile colonies (< 20 cm^2^, N = 1059) in white. Offspring assigned to at least one parent sampled within the study area, as revealed by parentage analysis, are shown in red (N = 216).

In total, 978 multilocus genotypes (MLGs) were detected in the entire population (from 3160 colonies genotyped) and 282 of them belonged to clonal lineages (many colonies of the same genotype) with GP values ranging from 3.1E-16 to 1.0E-06 for all loci combined. Overall, 84% of adults were clones indicating that asexual reproduction prevails for population renewal. Sixty-five percent of juveniles had at least one clone mate. In total, 677 juveniles were genetically identical to one of the sampled adults. Given the low *P*_(ID)_ value estimated (4.5E-08), these 677 juveniles (out of 1059) were removed from the pool of potential offspring as these clones are assumed to result from asexual reproduction through fragmentation. In order to perform parentage analysis, only the biggest clone within each MLG (i.e. the initial colony from which fragmentation most likely first occurred) was retained in our dataset. Following the removal of clonal replicates, 978 colonies with a unique MLG (genet level) remained to further assess sibship and parentage relationships among *M. platyphylla* colonies with a candidate pool of 603 parents and 375 offspring. Half of all sampled colonies were observed in the mid slope, where the proportion of juveniles was also much higher (56% of potential offspring) compared to what was found in the other surveyed habitats (25% in the upper slope and only 8% in the back reef) (Proportion test, *P* < 0.001, Table 1).

After the removal of clonal replicates, mean observed and expected heterozygosity were of 0.478 and 0.527, respectively, with 7.50 alleles per locus on average. The combined probability of exclusion, *P*_(EX)_, for the microsatellite marker panel was 0.94, revealing its high reliability for parentage assignments (Table S1).

### Parentage analysis and recruitment

Parentage analysis assigned 58% of all juvenile samples (216 of 375) to parents that were sampled within the study area (self-recruits, Fig. 2 and Table 1). Seventy-six percent of these self-recruits (165 of 216) were found within the mid slope, although the contribution from self-recruitment was significantly higher in the back reef habitat (Proportion test, *P* < 0.001, Table 1). Only 7% of self-recruits (25 of 375) had at least one parent located in one of the other habitats surveyed, suggesting more limited inter-habitat dispersal. Among the 603 parents sampled in the entire study area, only 96 of them contributed to self-recruitment in *M. platyphylla* population (Fig. S2). Forty-two parents were genetically unique, while the other 54 belonged to clonal lineages with a mean number of 12 ramets (potential parents) per MLG (range: 2–65). Clonal replicates for each of the 54 clonal parents were closely related in space with a mean distance of 15 m (± 28 SE; max = 226 m; median = 4 m). Of the 216 identified parent-offspring pairs, 203 were assigned to a single parent and only 13 were assigned to a pair of parents. Although the sex of individuals was not identified in this study, distance pairwise comparisons revealed that a mean distance of 314 m (± 281, SE) separated the two potential parents for the 13 parent-offspring pairs. Some of these parent pairs were very close to one another or very far from each other (4 to 892 m; median = 178 m) (Table S2). Furthermore, the mean pairwise genetic relatedness between parents contributing to self-recruitment (mean *r* = –0.011 ± 0.002 SE) was less than the average among all the potential parents surveyed in the study area (mean *r* = –0.002 ± 0.151 SE), indicating that inbreeding was limited.

While most local parents produced only one offspring each (51%), 32% of them produced three or more offspring within the study area. We identified four parents with more than five assigned offspring. One parent produced seven offspring and three other parents produced five offspring each within the study area (Fig. 3 and Fig. S3). Of these four families (one parent and several offspring), two were from clonal parents (Fig. 3C and Fig. S3). Clonal replicates and colony size of parents do not increase the number of offspring produced in the population (*r* = 0.067, *P* = 0.514 and *r* = 0.053, *P* = 0.605, respectively). Furthermore, the mapping of these families revealed that offspring settled parallel to the reef crest in alignment with the location of their parents (Fig. 3 and see Table S3 for details on each family).

**Figure 3.**
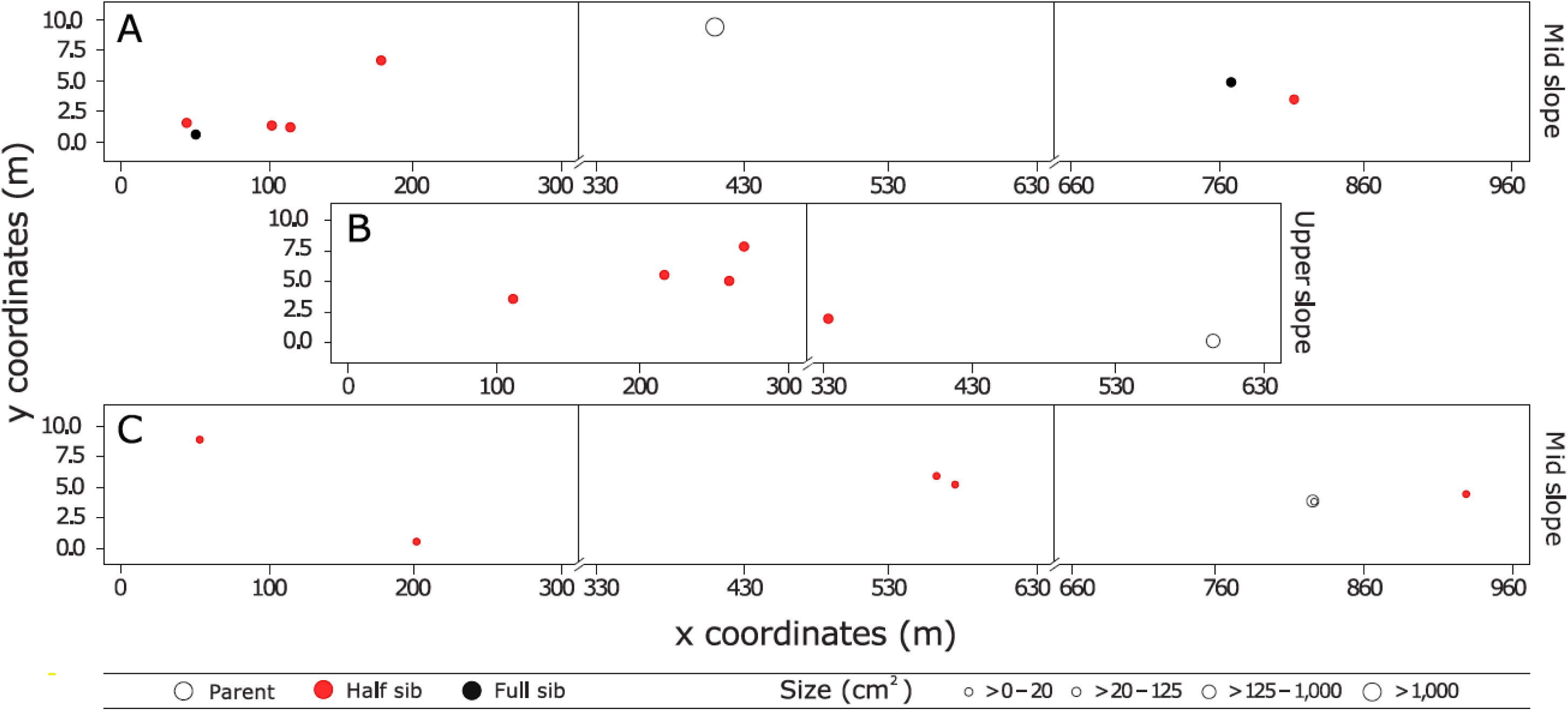
Dispersal patterns of three local families. Two are of parents with unique genotype (A) P1 with seven assigned offspring and (B) P2 with five assigned offspring, and one is of parents with clonal genotype (C) P3 with five assigned offspring. Dispersal estimates for parent-offspring relationship and mean dispersal distance among siblings are given in Table S3. One full sib relationship was found for P1 and an unsampled parent with a distance of 723.41 m between the two full siblings.

### Dispersal distances and sibling distribution patterns

Observed dispersal distances between offspring and parents ranged from 0.05 to 921 m (Table 2). Fifty percent of assigned offspring settled within 300 m of the parental sources with a significant gradual decrease in the proportions of offspring that settled at larger distances (r = −0986, *P* < 0.001, Fig. 4A). At a smaller spatial scale (100 m), our results demonstrate a higher proportion of offspring (34%) settling within the first 10 m from their parents (Proportion test, *P* < 0.01, Fig. 4B). Parent-offspring distances revealed no difference in the dispersal abilities of unique genotype parents *versus* clonal parents (Student’s test, *P* = 0.480, Table 2).

**Table 2.** Summary results of parentage analysis in *Millepora platyphylla* within the entire reef area (9000 m^2^). Characteristics of parents contributing to self-recruitment and comparisons between clonal and unique genotype parents are given: number of potential parents surveyed in the study area; number of parents assigned to an offspring within the surveyed area with their mean colony size (cm^2^), standard error (SE) and median; total number of assigned offspring and mean number of assigned offspring to one single parent with standard error (SE), maximum and median. All values are presented at the genet level (without clonal replicates). Estimates of offspring dispersal within the study area are shown: mean distance between parents and their assigned offspring with standard error (SE), minimum, maximum and median.

|  | Parent total | Parent clonal genotype | Parent unique genotype |
| --- | --- | --- | --- |
| # potential parents | 603 | 279 | 324 |
| # parents | 96 | 54 | 42 |
| Mean parent size (cm<sup>2</sup>) | 1 002.58 | 982.02 | 1 243.45 |
| SE | 2 980.19 | 2 950.73 | 3 336.22 |
| Median | 120.91 | 126.02 | 88.02 |
| # assigned offspring | 216 | 102 | 83 |
| Mean # assigned offspring per parent | 2.04 | 2.04 | 2.05 |
| SE | 1.30 | 1.30 | 1.45 |
| Max | 7 | 5 | 7 |
| Median | 1.50 | 2.00 | 1.00 |
| Mean dispersal distance parent-offspring (m) | 334.48 | 353.16 | 316.18 |
| SE | 231.47 | 261.99 | 219.74 |
| Min | 0.05 | 0.05 | 0.05 |
| Max | 921.37 | 921.37 | 870.25 |
| Median | 304.51 | 307.28 | 296.86 |

**Figure 4.**
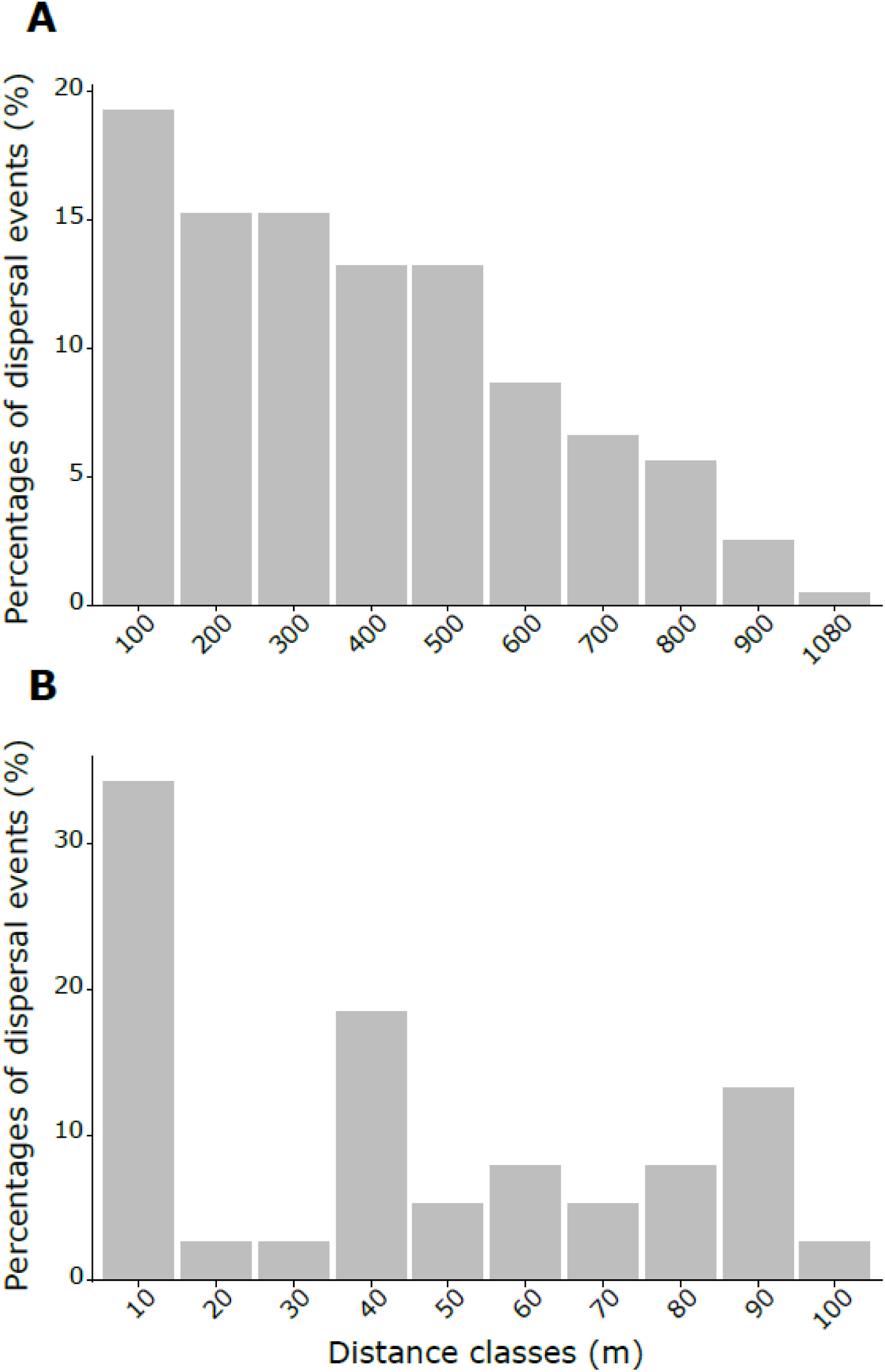
Dispersal distance distribution of parent-offspring pairs within the entire surveyed area. Observed dispersal distribution, determined using parentage analysis, is given by the percentages of dispersal events (N = 216) distributed among ten distance classes, assigned at 100 m each over the entire dispersal range (A) and assigned at 10 m each over the first hundred meters of the dispersal range (B).

Over the 375 offspring surveyed, 78 (21%) were involved in a full sibling relationship. Among these, 13 were assigned to local parent pairs, 40 to local single parents and 25 were from parents that were outside our field surveys. 132 offspring (35%) were also involved in a half sibling relationship from one local parent. No difference in the distribution of dispersal distances among full- and half-sibs was found (Chi square test, 1.96 > *P* > −1.96) and all siblings occurred within an aggregated pattern of distribution (within the first 100 m, Proportion test, *P* < 0.001, Fig. 5).

**Figure 5.**
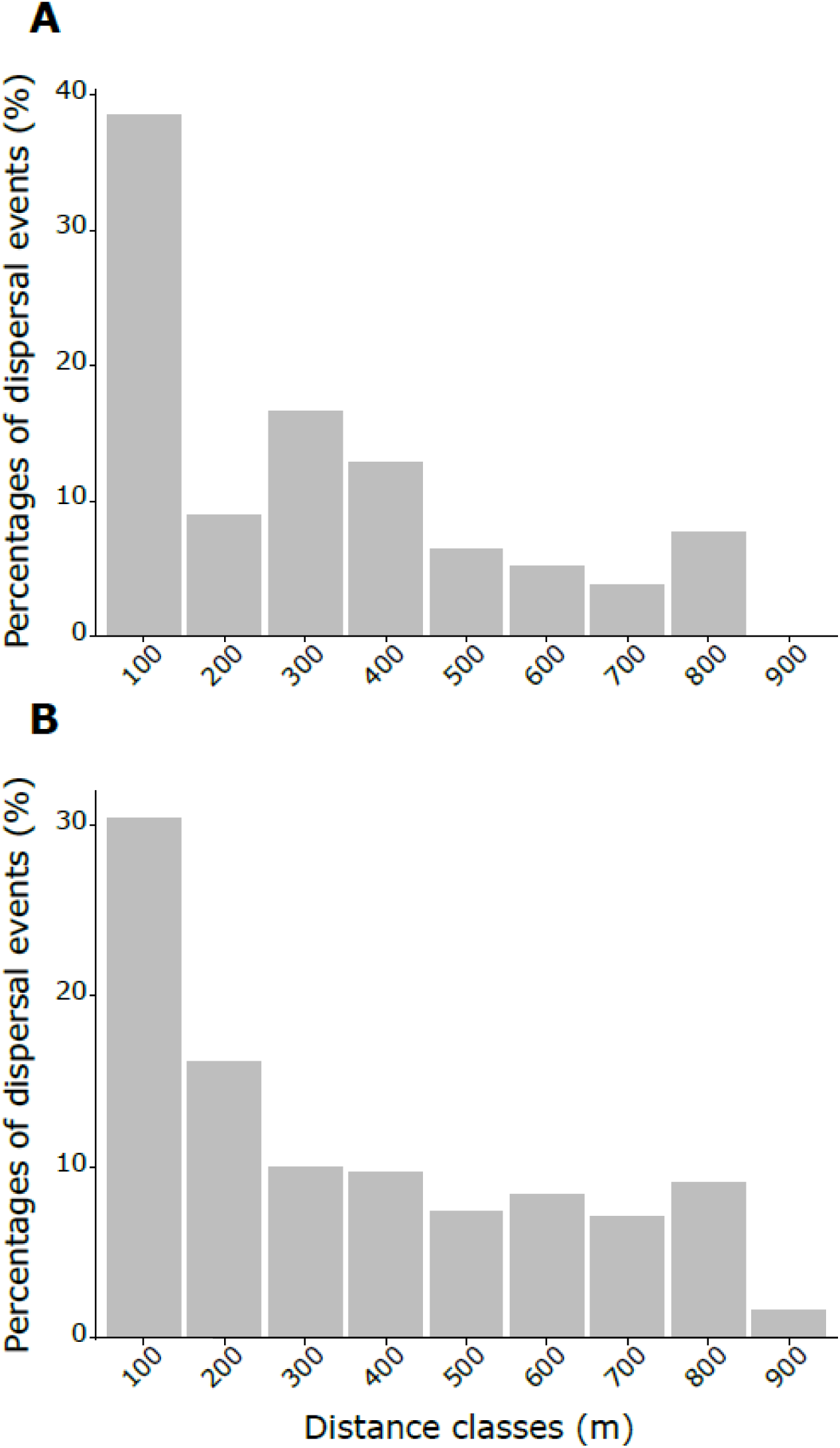
Dispersal distance distribution of siblings (full- and half-sibs) within the entire surveyed area. Observed dispersal distribution of full-sibs (A) and half-sibs (B), determined using parentage analysis, is given by the percentages of dispersal events (N = 78 for full siblings; N = 132 for half siblings) distributed among nine distance classes, assigned at 100 m each over the entire dispersal range.

## Discussion

By conducting genetic parentage analysis of 1059 juveniles and 2101 adults of the fire coral *Millepora platyphylla*, this study is one of the first to provide empirical estimates of dispersal in a broadcasting reef-builder. Our findings point to self-recruitment together with clonality as potential drivers of population recovery and persistence of *M. platyphylla* in Moorea. These results support earlier observations indicating that self-recruitment at the reef scale (or finer) governs the population replenishment of corals (Figueiredo et al., 2013; Gilmour et al., 2009; Underwood et al., 2018).

### High self-recruitment rate and limited dispersal within and among habitats

This study unveils a self-recruitment rate of 58% in *M. platyphylla* population, which is much higher than previous studies that focused on brooding species of gorgonian and scleractinian corals (∼25%, see Lasker et al., 2008; Warner et al. 2016). A high investment in self-seeding might well be an efficient process to sustain local populations in an isolated and fragmented reef system such as Moorea (as described in Gilmour et al., 2009; Shinzato et al., 2015; Tsounis & Edmunds, 2016; Underwood et al. 2018; Zeng et al., 2017). For *M. platyphylla,* we were expecting a lower contribution from self-recruitment because of its larval development mode (broadcast spawning). This reproductive strategy is often associated with high dispersal ability in natural populations of countless reef-building organisms (reviewed in Harrison, 2011). Consequently, such a high local recruitment does not exclude sporadic success of long distance dispersal events, significant in dispersing genetic variants.

Our results demonstrated that new recruits often settle within only few meters from their parents (less than 10 m). While very few studies have investigated the reproductive biology and early life stages of *Millepora* hydrocorals (reviewed in Lewis, 2006), our study revealed a limited dispersal ability of their sexual propagules (medusoids and planula larvae). Although medusoids can swim by pulsation of their bell, hydrocoral eggs are negatively buoyant and slowly sink after being released (Soong & Cho, 1998), and larvae crawl once they reach the reef substratum (Bourmaud et al., 2013). Such early life history traits are most likely rising opportunities for self-recruitment in populations of fire corals when sexual propagules are produced together with conditions that favour retention processes, settlement and post-settlement success. Still, before the larvae sink and crawl, there is extensive exposure to the currents in the water column during its early life history (medusoid and gamete release, fertilisation and larval development). The possibility of fire corals being able to produce brooding larvae cannot be excluded, although the confirmation of such a reproductive strategy in *M. platyphylla* is beyond the scope of this study. In addition, recruits with parents inhabiting distinct reef habitats that were separated by 40 to 160 m were recorded in small proportions (7%). Although limited, such dispersal events must rely on the passive dispersal of their sexual propagules through water circulation patterns. This possibility can easily be accomplished through wave-driven and cross-reef transports that characterise flow regimes on reefs located on the north shore of Moorea (Hench et al., 2008; Monismith & Herdman, 2013).

### Dispersal of early life stages with reef currents

In Moorea, alongshore and cross-reef transports are known to affect recruitment of larvae and population connectivity among habitats within a single reef, both in corals (Edmunds et al., 2010; Leichter et al., 2013, Tsounis & Edmunds, 2016) and fishes (Belgrade et al., 2012; Bernardi et al., 2012). Here, the low dispersal among habitats indicates that cross-reef transport, from the fore reef towards the lagoon, is not the major process driving the dispersal of offspring in *M. platyphylla*, although determinant for recruitment patterns in some scleractinian corals on Moorea’s reefs (Edmunds et al., 2010; Tsounis & Edmunds, 2016). However, these earlier interpretations were based on recruitment monitoring. Our genetic parentage data provide evidence that the dispersal of offspring crossing the reef matrix is a question warranting further investigations in scleractinian species. On the contrary, dispersal patterns within local families revealed that currents that run along the reef can help dispersing sexual propagules within the fore reef. This water mass circulating around Moorea’s reefs, together with early life history behaviour, results in the settlement of larvae parallel to the reef in alignment with the location of their parents. This dispersal pattern of sexual propagules is in contrast with the one reported for those that were produced asexually in the same fire coral population (Dubé et al., 2017b). In fact, asexual fragments were distributed perpendicularly to the reef crest, perfectly aligned with wave energy dispersal (see Fig. S2 for dispersal pattern of clones). A high proportion of juveniles was also reported on the mid slope, an exposed reef where wave energy is reduced (Hearn, 1999). As previously described in some Caribbean reefs (Paris & Cohen, 2004), water displacement decreases with increasing depth suggesting that larval behaviour, such as swimming or crawling, may enhance local recruitment in deeper waters within the fore reef. Overall, our results indicate that the distribution of fire corals in Moorea is strongly influenced by the dispersal of sexual propagules with along-reef currents, a process likely accentuated in deeper waters.

### Influence of gamete dispersal and fertilisation on sibling aggregations

Sibship analyses revealed that a large proportion of siblings complete their pelagic larval phase together. It is commonly assumed that medusoids, the early life stage during which a fire coral releases its gametes, facilitate fertilisation rates through synchronous spawning. This reproductive strategy enables gametes to aggregate at the water surface once they are released (Bourmaud et al., 2013; Soog & Cho, 1998) and most likely contributes to the sibling aggregation pattern observed in *Millepora* population. In many broadcast spawning species, the success of fertilisation is proximity dependent (Carlon, 1999; Doropoulos et al., 2018; Teo & Todd, 2018) with sperm dispersal identified as the limiting factor in numerous reef invertebrates, even at the scale of few meters (Coma & Lasker, 1997; Lasker et al., 2008; Pennington, 1985; Warner et al., 2016). In fire corals, we observed important variations in the geographical distance between identified pairs of parents (4 to 892 m with a mean of 314 m), highlighting a potential for fertilisation events between distant colonies. This result is not concordant with a recent study, where simulated intercolonial distances showed that fertilisation events rarely occur between coral colonies separated by more than 30 to 40 m (Teo & Todd, 2018). The authors argued that such a limited distance for successful mating from different coral colonies is mostly due to sperm dilution and insufficient mixing between gametes. In southern Japan, *M. platyphylla* males were found to release medusoids a few minutes earlier than females (Soog & Cho, 1998). This reproductive asynchrony may increase the probability of fertilisation success between distant colonies of fire corals.

### Impact of clonal reproduction on self-recruitment and dispersal

Even though adults were mostly clones, we found that genetically unique parents (only one ramet reproduces) contributed equally to self-recruitment than those of clonal genotypes (many ramets reproduce). Previous studies have shown that colonies that have suffered from stress due to fragmentation may further invest in growth rather than reproduction (Okubo et al., 2005; 2007). Since half of the clonal parents were smaller than 130 cm^2^, it is reasonable to assume that these fragments may preserve their energy to reach a larger size and increase their survival. Nevertheless, one could wonder if the propagation of clones increases the area over which sexual propagules are dispersed. In plants, the dispersal of seeds increases through clonal propagation and further reduces competition among siblings in the next generation (van Drunen et al., 2015). Our estimates for the dispersal distances were not higher for clonal parents suggesting that *Millepora* fragments have no better ability for the dispersal of their offspring. It has to be noted that asexual fragments were in close proximity to one another, which may further explain the similar dispersal extent of offspring among clonal and non-clonal parents.

Furthermore, most of the sampled juveniles were genetically identical to some parents. This result confirms that clonal aggregation of large colonies (i.e. when clones are distributed in patches) can increase local replenishment by supplying new recruits through their fragmentation (Highsmith, 1982). Despite the fact that large fragments are assumed to have a higher chance of survival (Lirman, 2000; Okubo et al., 2007), the large proportion of small fragments (< 20 cm^2^) observed in this study suggests that they can survive and effectively contribute to local sustainability. Still, these aggregations of clones combined with sibling aggregations and limited dispersal can increase inbreeding in the population due to cross fertilisation of genetically related neighbours. Despite all of these reproductive features, comparisons of genetic relatedness (all sampled parents *versus* those contributing to self-recruitment) revealed that mating between closely related adults is less likely to occur. This result suggests that the dispersal of sexual propagules, although limited, is enough to restrict population inbreeding. Furthermore, half of local parents relied on multiple breeding by reproducing with more than two other adult colonies within or outside the study area. As in numerous broadcast spawning marine invertebrates (Johnson & Yund, 2007; Lasker et al., 2008; Mokhtar-Jamaï et al., 2013; Warner et al., 2016), multiple mating may limit inbreeding in the population (Foerster et al., 2003) and increase the performance and survival of offspring by increasing the genetic diversity in the brood (McLeod & Marshall, 2009; Underwood et al., 2018). Dispersal limitations of both asexual and sexual propagules may represent an efficient reproductive strategy to sustain local population abundance and genetic diversity, especially at the margins of *M. platyphylla* range (Randall & Cheng, 1984).

## Conclusion

This study demonstrates that *M. platyphylla* is capable of producing enough sexual propagules to support self-seeding in Moorea’s populations, although they are predominantly sustained through asexual reproduction. The standing stock of genetic diversity in the population of fire corals at Moorea may also be eroded by its high level of clonality, hindering the potential for adaptation in the absence of input of exogenous genes (Uecker, 2017). The high self-recruitment rate observed in this study seems sufficient to provide the genetic diversity necessary for evolutionary adaptation. Such an asexual/sexual recruitment dynamics enables local sustainability and great opportunities to recover from major disturbances, which can occur frequently in coral reef ecosystems such as Moorea Island. This study confirms theoretical considerations claiming that self-recruitment is a key factor in stabilising population dynamics of reef organisms, such as fishes and reef-building corals (Hastings & Botsford, 2006; Richmond et al., 2018). Despite major differences in reproductive strategies, including clonal reproduction and planula larvae, reef fishes and fire corals have similar self-recruitment rates in Moorea (e.g., 30–60%, see Almany et al., 2007; Jones et al., 2005). Our study showed that the early life history of fire corals promotes the dispersal of sexual propagules locally as well as sibling aggregations. Strikingly, however, the dispersal distances seem extremely limited for a broadcasting species and raise uncertainties on how hydrocoral’s larvae can settle near their parental colonies. The dispersal assumption widely accepted for broadcasting corals make unlikely the possibility for a predominance of self-recruitment. Further investigations are thus needed to develop an understanding of the life cycle of fire corals and to determine early life history traits that are involved in the proliferation of locally produced larvae.

## Acknowledgement

We are grateful to Franck Lerouvreur and Marc Besson who helped with field surveys. Funding was provided to C.E.D. by the Fonds de Recherche du Québec – Nature et Technologies (graduate scholarship) and to E.B. by European Marie Curie Postdoctoral fellowship MC-CIG-618480.

## Data accessibility

Microsatellite loci for *M. platyphylla* are available in GenBank. All multilocus genotypes and sizes of reproductive colonies (including parents) and juveniles (including self-recruits) will be available upon acceptance.

## Author contributions

C.E.D. and S.P. designed research. C.E.D. and A.M. conducted field surveys and collected data. C.E.D. generated and analysed the data, and drafted the manuscript. C.E.D., E.B. and S.P. wrote the manuscript. S.P. acquired the funds for this study.

